# Automated STED nanoscopy for high-throughput imaging of cellular structures

**DOI:** 10.1101/2022.09.29.510126

**Authors:** Frank N. Mol, Rifka Vlijm

**Affiliations:** Molecular Biophysics, Zernike Institute for Advanced Materials, University of Groningen, Nijenborgh 4, 9747 AG Groningen, The Netherlands

## Abstract

STimulated Emission Depletion (STED) nanoscopy uniquely combines a high spatial resolution (20-50nm in cells) with relatively fast imaging (frame rate of ∼1-30Hz), straightforward sample preparation and direct image output (no postprocessing required). Although these characteristics in principle make STED very suitable for high-throughput imaging, only few steps towards automation have been made. Here, we have developed fully automated STED imaging, eliminating all manual steps including the selection and characterisation of the relevant (cellular) regions, sample focusing and positioning, and microscope adjustments. This automatic STED image acquisition increases the data output by roughly two orders of magnitude, resulting in a more efficient use of the high-end microscope, and the ability to detect and characterise objects that are only present in a small subset of the sample.

## Main

STimulated Emission Depletion (STED)^1^ nanoscopy is a light microscopy technique, which achieves high spatial resolutions (20-50nm) in cells^2^. STED thus can reveal important cellular structures that are too small to unravel using traditional methods like confocal and widefield microscopy, as these techniques are diffraction limited to resolutions >∼250 nm. In comparison to super-resolution PALM^3^ and STORM^4^ imaging, STED has the advantage of having a relatively high temporal resolution (frame rate up to ∼1-30Hz^5^). These short acquisition times, together with a straightforward sample preparation and direct imaging (no post-processing required), make STED a suitable technique for high-throughput imaging. However, the current imaging workflow requires time-consuming input by an expert user to manually find, select, characterise (e.g. 3D confocal overview) and STED image specific regions of interest (ROIs). Automating this workflow can exploit the full potential of STED. Besides a more efficient use of the high-end microscope and less manual labour for data acquisition, the probably most important benefits of automated high-throughput imaging are the decreased bias during data acquisition, and the significantly improved statistics and thus increased reliability of the results. Furthermore, automation increases the opportunity to detect transient structures without (chemically) altering cellular processes. Thus, to overcome the current low-throughput, there is a demand for unsupervised fully-automated STED imaging, which is not met up to date.

Thus far, automated high-throughput imaging has been successfully developed for several lower-resolution microscope techniques (e.g. Micropilot^6^), including methods that adjust microscope settings (Micromator^7^). Although similar methods do not yet exist for STED nanoscopy, an important step has been made by enabling microscope control through Python code (ImSwitch^8^ for home-built systems and Imspector^9^ for commercial Abberior microscopes). This basis allowed for the imaging of regions larger than the typical field of view (FOV) by STED^10^, and the implementation of event-triggered STED nanoscopy^5^. While these important developments enable to image structures larger than the typical FOV or at a relevant point in time at a specific location, they do not yet enable automated high-throughput imaging of large numbers of structures.

In this work, we present a method for unsupervised, fully-automated detection and imaging of objects using STED nanoscopy. The protocol starts by scanning a large sample area using fast and low-resolution imaging to localise the ROIs that contain the objects which should be measured by STED. These ROIs are classified, after which a selection (either random or based on predefined features) is further characterised at higher resolution, and finally imaged using STED nanoscopy. To exemplify this approach, we have used automated STED nanoscopy to image ESCRT-III proteins in unsynchronised wild type populations of fixed *Sulfolobus acidocaldarius* cells, with the goal of characterising the structure of division rings at different stages in the division process.

## Results

The automation that we have developed enables high-throughput STED imaging by detecting objects in a large sample area (many FOVs, not limited), and applying STED microscopy in an unsupervised manner. This automation was achieved through dedicated python software which enabled the control of microscope settings and stage positions, and further included image analysis and decision making. Our workflow consists of several sequential steps: (i) (low resolution) sample overview, (ii) segmentation, (iii) classification, (iv) selection, (v) characterisation and (vi) STED measurement (Fig. 1). Each of these steps is discussed in further detail below.

**Figure 1.**
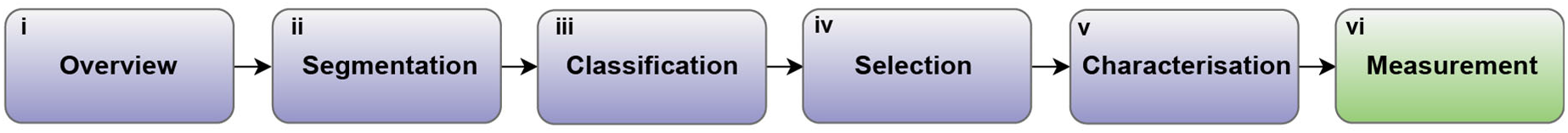
Workflow for automated STED measurements. (i) In the first step of our automated workflow low resolution overview images are acquired. Multiple FOVs are measured to obtain a large scanned area of the sample. (ii) The second step is the segmentation of objects of interest as separate objects in the overview measurements. The centre positions of these objects are translated to the stage coordinates of the microscope. (iii) Next, all segmented objects are classified based on the joint presence of objects and object features such as morphology and intensity. (iv) The objects to measure using STED are selected from the overall list of objects. Either a subset (based on characteristics as determined in the classification step), or a random subset of the complete set can be selected. (v) Of each selected object, a confocal z-stack is measured of all fluorescent labels in the sample. (vi) In the final step, the object is measured in STED mode. The stage positions for this scan are obtained from x,y,z-fitting of the z-stack images. The FOV is minimised for minimum impact on the cells, minimum drift, and maximum imaging speed of the consecutive STED image.

### Overview

The first step of the automation workflow is obtaining an overview of the sample placed on the microscope. This overview is used to characterise the sample and identify objects. The imaging modality and parameters used for the overview images should provide sufficient detail to identify possible regions of interest, and at the same time minimise the imaging time and light dose (to reduce phototoxicity and photobleaching). It is important to optimise the pixel size (as large as possible) and dwell times (as short as possible) to minimise the acquisition time and imaging impact, while still enabling segmentation. If merely the locations of prominent features (e.g. cell locations) need to be found, a lower magnification objective could be used and/or epifluorescence illumination in combination with camera based detection. A simple coordinate transformation between this imaging modality and the subsequent confocal/STED imaging would suffice to obtain correct microscope stage positions for the latter. In case smaller features (e.g. subcellular structures) need to be detected for further decision making, these overviews should be obtained by confocal imaging. In our example, we used confocal microscopy. As a single FOV in our microscope measures 80×80μm^2^, tiling was used to obtain a large overview. In our tiling approach, multiple adjacent FOVs were obtained by automatically moving the microscope stage by 80 μm, after which the images are stitched together. The quality of the stitching can be improved by having a slight (e.g. up to 10%) overlap of neighbouring FOVs, but for merely detecting regions of interest, moving the stage a full FOV is often desired in view of photodamage and imaging speed. As an alternative to tiling, non-overlapping regions (randomly chosen) from a much larger sample area could be imaged. The benefit of this approach is that in the same acquisition time, a broader, less region-dependent sample overview can be obtained. To extend the axial information, a z-stack can be recorded. Again, in view of speed and photodamage, the height increment between z-slices should be chosen at the maximum distance that still enables sufficient object detection. In our example, we obtained a z-stack consisting of three slices.

With regard to the applied illumination and possible excitation/detection multiplexing scheme, the minimum number of colour channels should be used that still enables object identification. In the here presented example study (see Fig. 2), we imaged confocal overviews in 4-colours, which were frame-multiplexed (individual frames for each if the 405nm, 488nm, 561nm and 640nm excitation lasers). Frame-multiplexing enabled to trace back the fluorescence signal to the individual fluorescent labels (Hoechst, Alexa Fluor 488, Abberior STAR 580 and Abberior STAR 635). All overview images combined should at least contain enough objects to fill the desired experiment runtime with STED imaging, but ideally more overview images should be collected to enable data selection prior to STED imaging. In our example, 5×5 FOVs (400×400µm^2^) contained sufficient objects for an experiment runtime of ∼3h. For an overnight run of ∼12h, the overview size was set to 10×10 FOVs (800×800µm^2^). The acquisition time of an overview image depends on the sample brightness (brighter samples require less dwell time) and the number of frames per FOV (e.g. slices in a z-stacks, multiplexing instead of collecting all colour channels in a single scan, see Table 1), ranging from 1s (1 slice) up to 25s (multi-colour z-stack). It furthermore takes ∼0.3s to move of the sample from one overview position to the next, using a standard Olympus motorised stage. We repeat the positioning of the stage three times to improve the precision of the stage position, resulting in an overall movement time of ∼0.5s (the last two positionings require minimal stage movement and are thus faster). Storing the overview images (four z-stacks, consisting of 3×400×400 pixels) took around 16ms.

**Table 1.** typical acquisition times per FOV. Additionally ∼0.3-0.5s are required for the stage movement and ∼16ms for image storage.

| microscopy type | multiplex | acquisition time | field of view |
| --- | --- | --- | --- |
| **confocal** | off (all colours simultaneous) | 1-2s | $80 \times 80 \mu\text{m}$ |
| **confocal** | 4 colour sequence | 4-8s | $80 \times 80 \mu\text{m}$ |
| **widefield** | off (all colours simultaneous) | 1-2s | $103.5 \times 77.64 \mu\text{m}$ |

**Figure 2.**
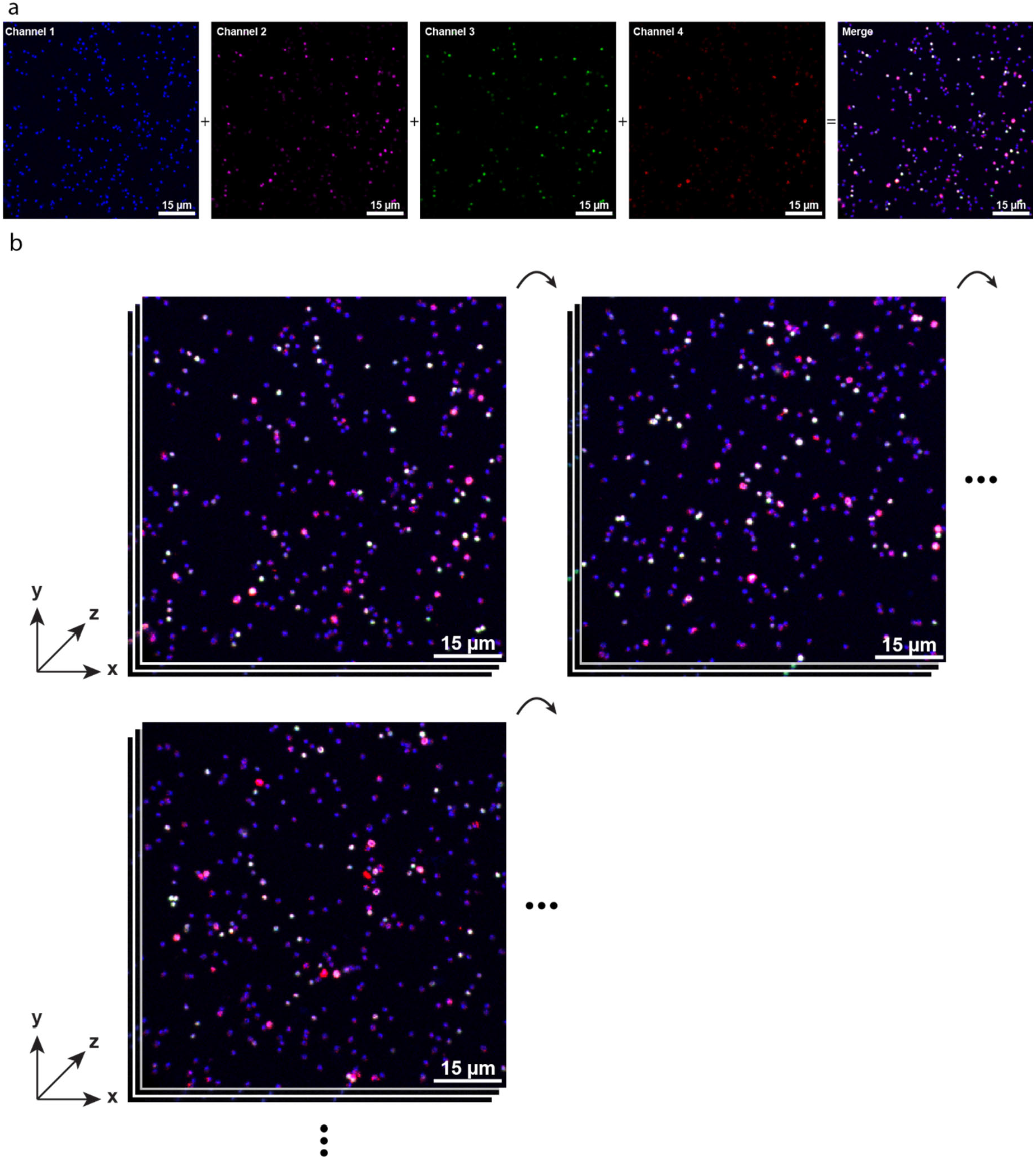
Overview image acquisition of images, in which the DNA and the different ESCRT-III proteins in *Sulfolobus acidocaldarius* cells have been labelled with Hoechst (DNA), Alexa Fluor 488, Abberior STAR 580 and Abberior STAR 635 respectively. (a) From each FOV (80×80µm^2^), all four labels were separately imaged: DNA (blue), CdvB2 (magenta), CdvB1 (green) and CdvB2 (red). (b) At every stage position, a z-stack consisting of three slices was created (120nm spaced). Tiling was used to obtain a large summary of the sample. 5×5 FOVs (400×400µm^2^) and 10×10 FOVs (800×800µm^2^) are obtained for short runs (∼3h) and long runs (∼12h), respectively. The pixel size was 200×200nm^2^ to minimise photobleaching.

**Figure 3.**
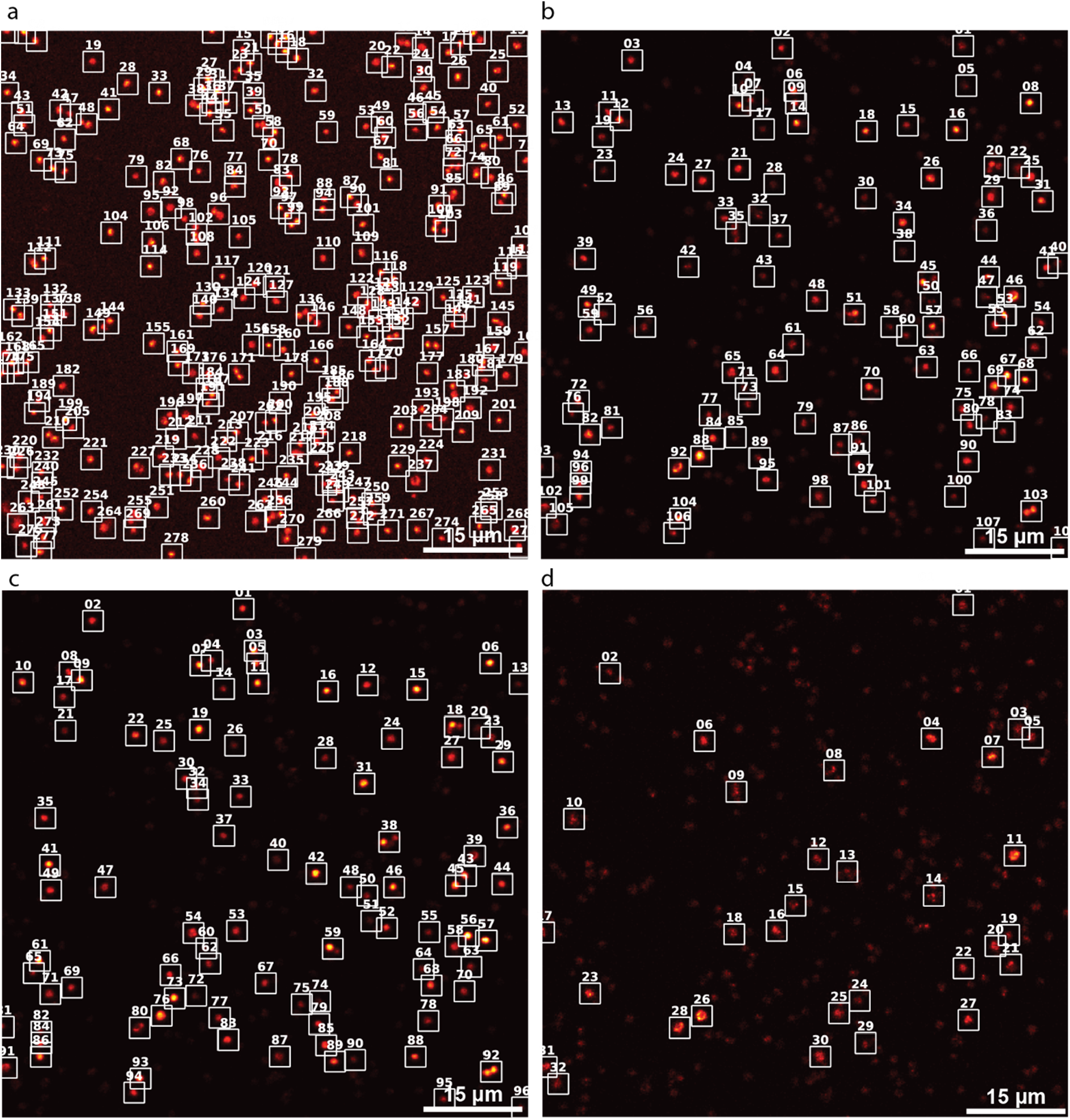
Segmentation performed on every label of the overview images to obtain the objects and their corresponding coordinates. The presented segmentation maps correspond to (a) DNA labelled with Hoechst, excited with a 405nm laser, (b) CdvB2 labelled with Alexa Fluor 488, excited with a 488nm laser, (c) CdvB1 labelled with Abberior STAR 580, excited with a 561nm laser and CdvB labelled with Abberior STAR 635, excited with a 640 laser. The overview images are averaged in the z-dimension (the mean of the three slices z-stack) to segment objects, which lie in different focal planes.

### Segmentation

After obtaining the overview images, the labelled objects of interest (and their corresponding stage positions) are extracted through segmentation. In our example study, already a simple intensity thresholding proved sufficient to discriminate objects from background and thereby allowed to segment the ROIs. To reduce the impact of noise, the overview images are smoothed using a Gaussian Blur before thresholding. By adapting the thresholding, bright objects with high intensity as well as fainter objects can be selectively identified. After thresholding, a combination of opening and closing sequences is applied.

To determine the quality of this approach, we analysed the segmentation accuracy for overview images of fixed *Sulfolobus acidocaldarius* samples, in which several cell division proteins (CdvB, CdvB1 and CdvB2) and DNA were fluorescently labelled. We manually annotated segmented overview images and verified for each of these labels, which of the segmented objects should not have been segmented, and which of the objects had not been selected by the segmentation algorithm. Using 25 FOVs (∼28k cells), the annotated segmentation maps resulted in a precision (correctly identified objects among all automatically segmented objects) between 0.978 and 0.999 (Supplementary 1), and a recall (fraction of relevant objects that was segmented) between 0.731 and 0.901 (Supplementary 1) for the labelled ESCRT-III proteins. When determining optimal segmentation parameters, we optimised for better precision instead of better recall to reduce the number of STED images containing no object of interest while accepting to miss a few possible STED imaging locations, since the output was mostly limited by the characterisation and (STED) measurement time. The thresholding can be further optimised for every new type of sample. However, in general, most samples primarily consist of (lower intensity) background, and our approach is thus widely applicable. In case a sample requires more sophisticated segmentation (e.g. the sample does not contain many objects, or the objects of interest display less discernible features), deep learning applications^11,12^ can be implemented in a modular fashion. In our example, we have collected segmentation data of four labels in a single sample. For each of these colours, a list of segmented ROIs was created. In preparation for the consecutive measurement steps, which involve microscope stage movements, the pixel-coordinates of each of the centres of the segmented ROIs were transformed to stage-coordinates and stored in lists. Here, we have shown that this relatively straightforward segmentation proved to be efficient (see precision and recall, Supplementary 1) on fluorescently labelled ESCRT-III proteins and DNA in archaea (Supplementary Fig. 1).

### Classification

An important benefit of automated imaging is the opportunity to quantify the sample composition. From the segmentation data, many features can be extracted. If the chosen labelling allows to identify individual cells (e.g. via DNA staining), all segmentation layers can be correlated to the identified cell positions. This results in a quantified sample composition with regard to for example the morphology, the combination of proteins present and their intensities, and the relative distances between proteins. For each of the segmented archaea cells (based on the location of the DNA signal) in our example study, we quantified the shape, size and the mean intensity of each labelled cell division protein (CdvB, CdvB1, and CdvB2) (Fig. 4). Furthermore, the combination of the different protein labels present in each cell was evaluated (Supplementary Fig. 3) and stored together with these quantifications and the corresponding stage coordinates in a list. In the current study, seven different combinations of present proteins are possible, ranging from the detection of one of the three proteins to the joint presence of all three labelled proteins.

**Figure 4.**
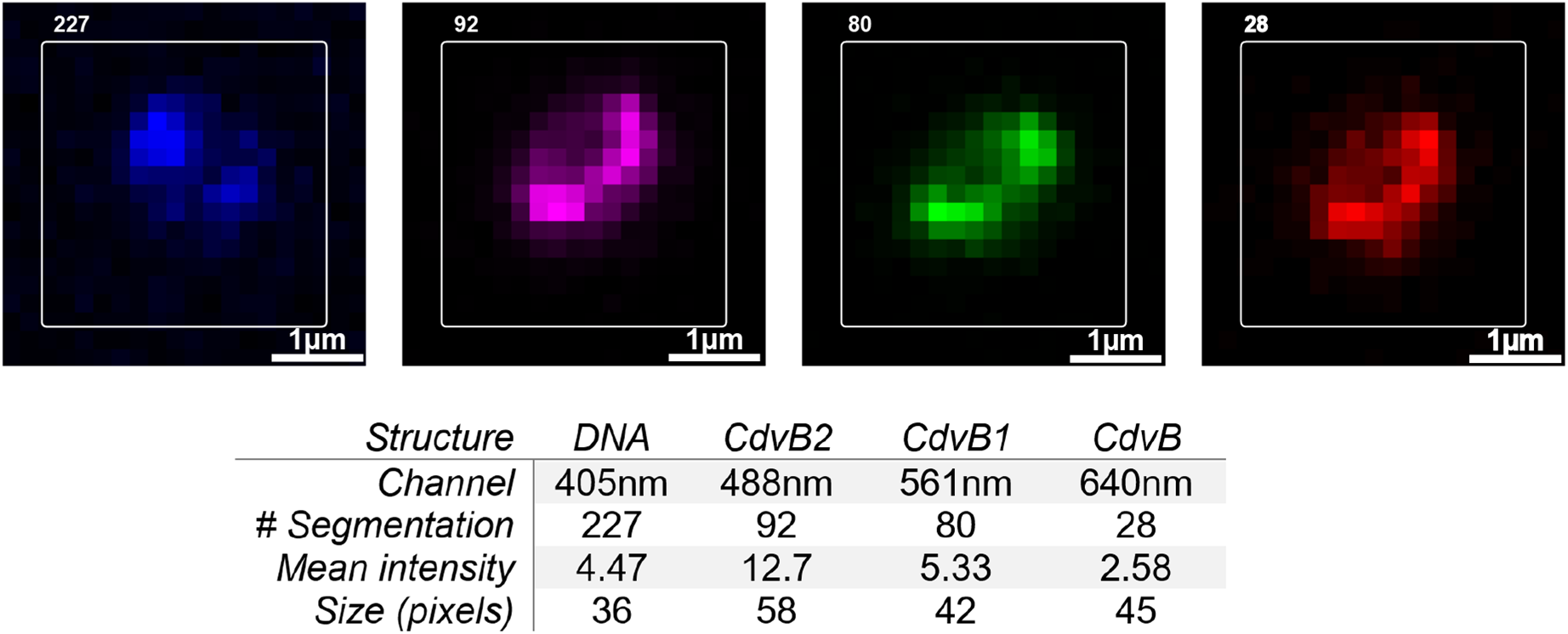
Classification of the segmented ROIs. The presented example shows joint presence of all proteins (CdvB, CdvB1 and CdvB2). The individual numbers correspond to the segmentation number in the individual channels, which can be used to locate a measured object in the overview images. Furthermore, for each channel, the mean intensity per slice of z-stack is stored, which can be used to determine the z-position, in which the object is in focus. The intensity, next to information on the morphology, is also stored to be later used in the selection step of the workflow. The morphology information that we used in the guiding example is the size of the segmented object.

### Selection

Traditionally, a microscopist selects the cells/objects to be imaged based on experience and/or expectation of what a good cell/object should look like, without taking any quantitative selection criteria as basis. Even if deliberately aiming for a random and unbiased object selection, by nature, the human eye is not very well suited for this task. Uniquely, automated imaging allows to select a subset of objects based on well-defined quantitative object features, after a full characterisation of all objects in a large sample area is achieved,

Three approaches can be applied in the object selection. The most straight forward one is to obtain STED images of all objects within the sample that qualify for a defined combination of features (e.g. protein A and B should be present, proteins C should be absent). However, measuring the complete sample can take a large amount of time to measure. Hence, choosing a subset objects that qualified for acquisition often is preferred. A selection of objects can be made either at random to obtain a pool of STED images representing the sample composition, or based on specific object characteristics. For example, the selection can depend on features such as the object’s size or its intensity. Quantitative rules then determine if an object lies within the subset of object to be imaged by STED. There are many different ways, in which these rules can be implemented (e.g. brightest x%, or equal ratios of the least bright 33%, medium bright 33% and brightest 33%). Key feature is the defined selection process leading to the final dataset, which allows the scientific community to directly evaluate the applied criteria. With progressing insight into the objects studied, a revision of these criteria can be made and other subsets of the data satisfying adapted selection criteria can be recorded. Also, even if only a small subset is selected for the final STED measurements (Fig. 5), knowledge on the sample composition as obtained in the previous low-resolution steps is still very valuable. In the present study on archaea, we selected a random subset for each given label combination (Supplementary Fig. 3). Furthermore, a subset was selected that represented the highest protein concentrations within the selected ROIs (highest average photon count in the object). We determined the number of objects to be imaged based on the intended runtime of the complete automation workflow. Note that the random subset was selected first and the high intensity subset was picked out of the remainder of objects to obtain a truly random dataset reflecting the sample composition.

**Figure 5.**
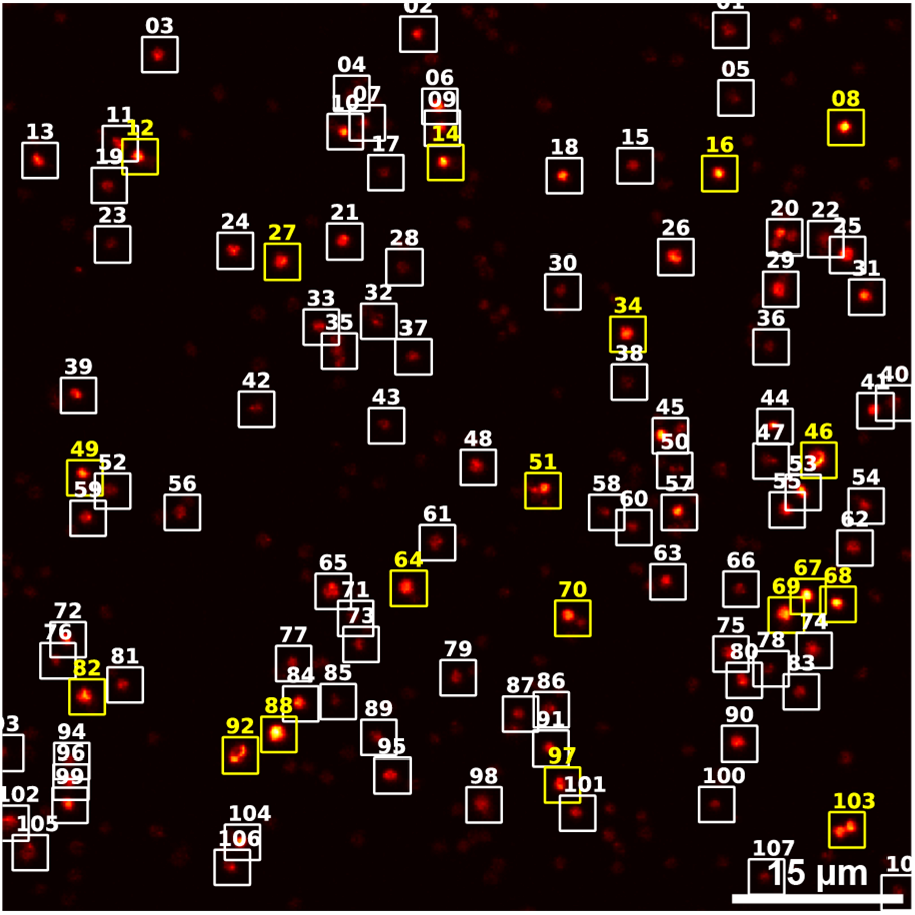
Selection of the segmented and classified objects. To obtain sample representative measurements, a random subset of the segmented ROIs can be selected. However, instead, the quantifications from the classification step can be used to select a specific subset of the identified ROIs. The presented example is a segmentation map of the 488nm channel consisting of labelled CdvB2. The FOV is 80×80µm^2^. The yellow boxes show which of the objects are selected to be measured using STED.

### Characterisation

To characterise the selected objects in more detail, the stage is moved to the coordinates of each selected object sequentially. At a given stage position, a confocal z-stack is recorded to perform a 3D characterisation. This z-stack serves two purposes. First, it shows in greater detail the 3D distribution of all labelled proteins in the specific cell, as often more than just the proteins, which should be imaged by STED, are labelled (e.g. for our archaea samples, DNA is labelled additionally to the three cell division proteins, Fig. 6). Hence, important context for the final STED image of the selected object can be obtained in this step. Second, with the z-stack, the precise lateral and axial position of the protein(s) of interest is obtained through an x,y,z-fit, enabling optimal focusing in the final STED measurement. Instead of applying a fit to the z-position, selecting the slice which has the maximum mean intensity often also suffices. It is worth to evaluate the optimal pixel size for the confocal 3D stack. As confocal imaging does not reach the resolution obtained in the final STED image, an intermediate pixel size of 80×80nm^2^ is often reasonable to choose. If the information in this 3D stack does not require this precision, selecting a larger pixel size will limit possible phototoxicity or photobleaching and speed up the runtime. The same reasoning should be applied to the choice of the z-stack spacing. Here, an initial choice of 120nm or more is reasonable.

**Figure 6.**
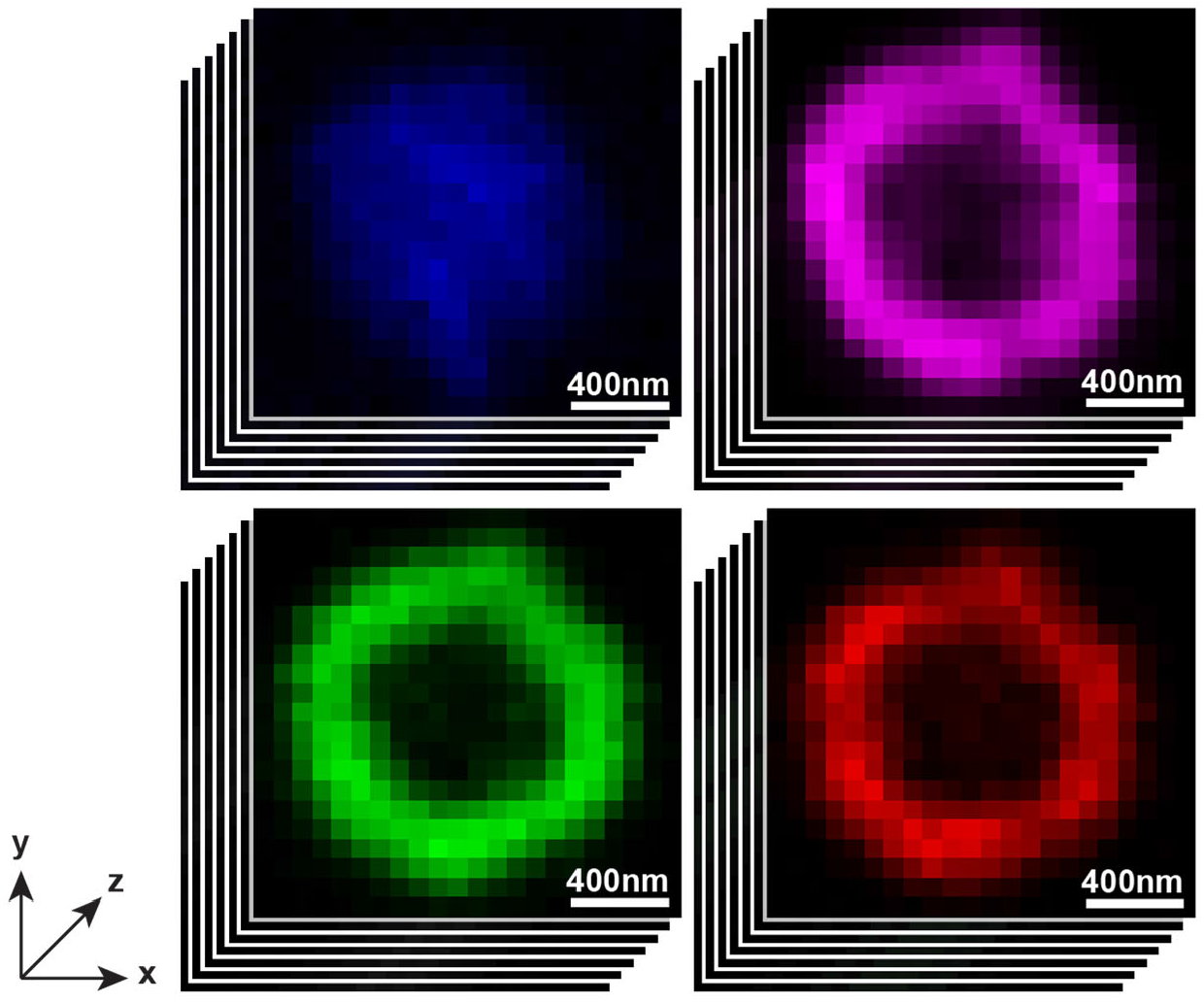
Characterisation of an individually selected object of interest. performed by a confocal z-stack, consisting of twelve slices (spaced 120nm), a FOV of 2.96×2.96µm^2^ with 80×80nm^2^ pixels. The z-stack enables to visualise the structure and its environment in a 3D context. Furthermore, an x,y-fit is used to find the optimal lateral stage position for the final STED image. An additional z-fit or just the selection of the brightest slice of the stack enable to position the stage in the axial direction for the subsequent STED image. The z-stack consists of DNA labelled with Hoechst (blue), CdvB2 labelled with Alexa Fluor 488 (magenta), CdvB1 labelled with Abberior STAR 580 (green) and CdvB labelled with Abberior STAR 635 (red).

### Measurement

In the final step of our workflow, the selected objects are measured in STED mode. As in the previous characterisation step the x,y,z-positions were fitted, optimal focusing for STED imaging is set in an unsupervised manner. The lateral positions as previously identified in combination with knowledge on the object size allows to image the smallest possible region with STED. As STED is a scanning technique, this approach reduces the time to perform a measurement, resulting in faster data acquisition, less sample drift during the measurement, and less phototoxicity when performing live cell measurements. To further minimise photodamage, adapted acquisition schemes such as ResCUE^13^ and DyMIN^14^ can be applied. In our study, all selected objects were measured within a FOV of 2.5×2.5µm^2^, using a pixel size of 17×17nm^2^, while we obtained a resolution of around 30nm^15^ (Fig. 7). With our chosen measurement conditions, the detailed confocal z-stack (important to give context to the detected objects measured by STED), the analysis, and the sequential dual colour STED measurement all together had a total runtime of ∼68s per object.

**Figure 7.**
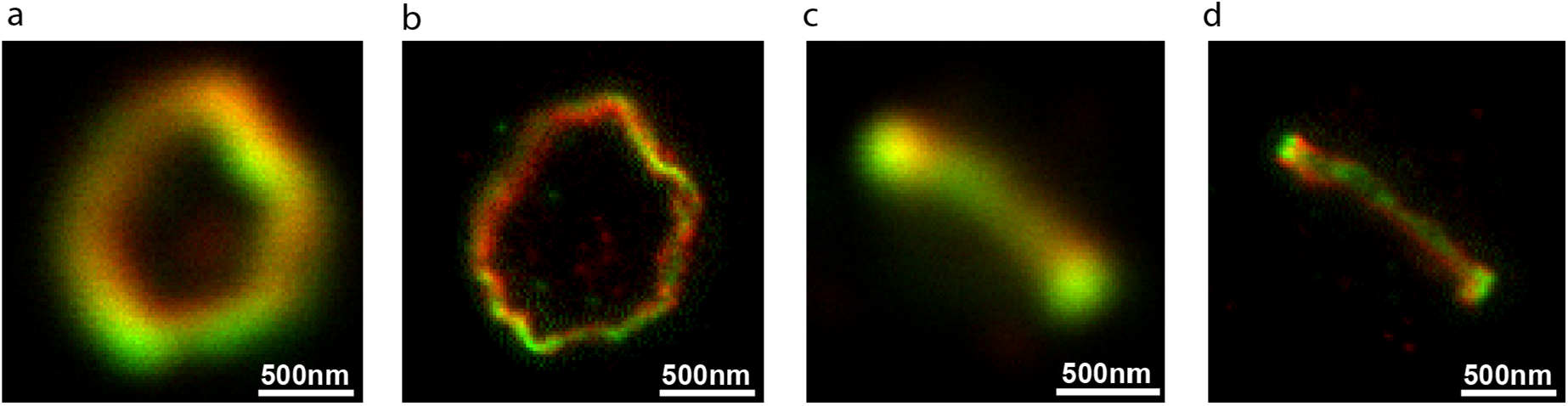
Acquisition of STED images. Example images of confocal and corresponding STED images of objects found in the previous sample characterisation steps. (a-d) ESCRT-III protein ring consisting of CdvB1 labelled with Abberior STAR 640 (red) and CdvB2 labelled with Abberior STAR 580 (green). (a-b) Face-on representation of a protein ring, measured using (a) confocal and (b) STED. (c-d) Perpendicular representation of a protein ring, measured using (c) confocal and (d) STED. The pixel size in the STED images is 17×17nm^2^. The presented STED images have a resolution of around 30nm.

For the guiding example, we studied three different sample conditions in triplicates and with reversed labelling as a control (i.e. swapping labels on target proteins). We thus measured 18 different samples and obtained a total of 5,586 dual colour STED images at 30nm resolution. All these images are supplemented with a confocal z-stack revealing up to three proteins and DNA. Furthermore, by the automatic bookkeeping, the imaged object can be found back in the segmentation maps for each of the four channels, allowing to analyse possible relations between the measured objects and the sample (e.g. sample density or spatial clustering of specific events). In a similar project we have measured peroxisome proteins in yeast. For this sample we found the same segmentation strategy to be again highly efficient (Supplementary Fig. 2), enabling us to analyse STED imaged objects of even more (8,063) individual cells.

## Discussion and conclusion

Since its invention^1^, great progress has been made on all aspects of STED nanoscopy, enabling multi-colour imaging with 20-50nm resolution in (living) cells. STED even reaches sufficient resolution to detect the spatial separation between the N and C termini of the same protein^16^. Developments of the microscope technique itself have brought important optimizations improving the acquisition time^17^, and decreasing the imaging impact^18-21^ and imprecisions caused by for example aberrations^22,23^. These developments have made STED nanoscopy a widely used method to visualise cellular processes. Unique in comparison with other methods that obtain similar resolutions^24-26^ is the very short acquisition and image processing time of STED. Although some techniques obtain a better spatial resolution (e.g. MINFLUX^27^) or temporal resolution (fast camera-based imaging^28^), STED uniquely combines high spatial (20-50nm^2^) and temporal (framerate of 1-30Hz^5^) resolutions in live cell imaging. Unfortunately, even though STED imaging itself can take significantly less than a second, studies using this method are typically still based on few tens to few hundreds of acquired images, since prior to STED imaging, time consuming steps in object selection, characterisation and focussing are required. Another drawback of the characterisation steps is the potentially induced phototoxicity and photobleaching of the fluorescent labels. To achieve minimal photodamage, each characterisation step should be executed with different microscope settings (including changing between widefield, confocal and STED nanoscopy and acquisition settings like illumination intensities, pixel sizes, pinhole diameters, dwell times and microscope stage positions). Lacking any automation in this process so far, all current workflows demand manual microscope adjustments, which are strongly dependent on the user response time. The limiting factor in data acquisition is therefore the manual steps prior to STED imaging, and not the time it takes to acquire the actual STED image.

In this work, we have achieved completely automated STED imaging of labelled DNA and ESCRT-III proteins in archaea cells, without any user input after placing the sample on the microscope stage. We have eliminated all time-consuming manual elements, including the selection of data points, characterisation of the relevant (cellular) regions, stage positioning and focussing, and microscope adjustments. This automated image acquisition in STED microscopy increases the data output by roughly two orders of magnitude resulting in a more efficient use of the high-end microscope. This increase in the dataset size leads to better statistics and hence more reliable conclusions. Furthermore, as opposed to the conventional procedure, in which a microscopist manually selects objects to be imaged, our selection parameters are quantifiable and made explicit, and thereby can lead to a reduction in an implicit selection bias by the microscopist^29^. The stored overview images and segmentation information furthermore allow for post-analysis of the sample composition. Finally, a major advantage of using the proposed automation method is the ability to acquire data from rare states, such as late division rings in archaea. Without automation, these rare states are hard to find manually. A solution, which is generally applied to image such rare states, is to biochemically force cells into those specific states. Uniquely, our automation method now enables the imaging of these rare states in unmodified, asynchronous populations of wild type cells.

We have demonstrated the reliability of our method by imaging 5,586 dual colour STED images at 30nm resolution in wild type samples of archaea, requiring the selection of only the dividing cells. Our results highlight that the automation made an incredibly large study possible with a feasible measurement time and with very limited handling time at the microscope.

## Materials and Methods

### Microscopy setup

The microscope used for performing the automatic measurements was an Abberior Expert Line STED microscope. The used objective was an 100x immersion oil objective (Olympus Objective UPlanApo 100x/1.40 Oil). Prior to imaging, the lasers were aligned using 0.1µm TetraSpeck™ (Invitrogen, T7279) Beads. The excitation was done by a continuous wave laser of 405nm wavelength (2mW at laser head), a 40MHz pulsed laser of 488nm wavelength (220µW at laser head), a 40MHz pulsed laser of 561 wavelength (200µW at laser head) and a 40MHz pulsed laser of 640nm (1mW laser at head). The depletion was done by a 40MHz pulsed laser with wavelength 775nm (3.2W at laser head). Avalanche Photodetectors were used for each laser channel.

### Imaging parameters

The overview images were created as a confocal z-stack consisting of three slices (spaced 350nm) each had a FOV of 80×80µm^2^. The pixel size used was 200×200nm^2^. The pinhole was set to 3.0 A.U.. The excitation laser and detection window settings were set as follows: the 405nm laser at 8% (160µW at laser head) and a detection window of 415-478nm. The 488nm laser at 5% (11µW at laser head) and a detection window of 505-550nm. The 561nm laser at 10% (20µW at laser head) and a detection window of 575-630nm, and the 640nm laser to 0.1% (1µW at laser head) and a detection window of 650-700nm.

A confocal z-stack consisting of twelve slices (spaced 120nm) was acquired to characterise the relevant objects, with a FOV set to 2.96µm^2^, a pixel size of 80×80nm^2^, and a pinhole of 0.8 A.U.. The excitation laser and detection window settings were set as follows: The 405nm laser at 15% (300µW at laser head) and a detection window of 415-478nm. The 488nm laser at 15% (33µW at laser head) and a detection window of 495-550nm. The 561nm laser at 20% (40µW at laser head) and a detection window of 575-630nm, and the 640nm laser at 0.4% (4µW at laser head) and a detection window of 650-763nm.

The STED measurements images were created with a FOV set to 2.5×2.5µm^2^ with a pixel size of 17×17nm^2^ and a pinhole set to 0.8 A.U.. To reduce the photobleaching, ResCUE^13^ and DyMIN^14^ were used. The settings of the excitation and depletion lasers were sample dependent. Six different combinations of samples were measured. The corresponding settings are presented in Table 2.

**Table 2:** Excitation and depletion settings for the STED images.

| 640nm channel | 561nm channel | Line steps |  |  | Intensity Confocal laser (%) |  |  | Intensity STED doughnut (%) |  |
| --- | --- | --- | --- | --- | --- | --- | --- | --- | --- |
|  |  | confocal | Low int. STED | High int. STED | confocal | Low int. STED | High int. STED | Low int. STED | High int. STED |
| CdvB |  | 5 | 6 | 19 | 0.4 | 0.5 | 1.1 | 4.6 | 45 |
|  | CdvB | 2 | 8 | 8 | 7.8 | 13 | 20 | 15 | 90 |
| CdvB1 |  | 3 | 5 | 18 | 0.2 | 0.45 | 1.1 | 3.6 | 45 |
|  | CdvB1 | 1 | 4 | 5 | 3 | 4 | 15 | 12 | 90 |
| CdvB2 |  | 3 | 5 | 18 | 0.2 | 0.45 | 1.1 | 3.6 | 45 |
|  | CdvB2 | 2 | 10 | 8 | 17 | 20 | 25 | 15 | 90 |

### Sample preparation and cell lines

The archaea samples used for imaging in this work were prepared as described in Hurtig et al.^15^. DNA was labelled with Hoechst and imaged in the 405nm channel. The ESCRT-III proteins (CdvB, CdvB1 and CdvB2) were labelled with Alexa Fluor 488, Abberior STAR 580 and Abberior STAR 635 and imaged in the remaining channels (488nm, 580nm and 640nm, respectively).

Samples of *Hansenula polymorpha yeast* cells were prepared similar as described in ref ^30^. First, they were grown overnight at 37 °C, after which they were diluted to an OD of 0.1 in mineral medium (MM)^31^ supplemented with 0.5% glucose as carbon source. Next, they were grown for 6h on MM supplemented with 0.5% methanol as carbon source to obtain an OD of 0.5. For the experiments 5 OD units were harvested. At this point, the cells were fixed (30 min, 2% formaldehyde in 500 ul PBS on ice in dark), and stained for 1h with 1 μM SiR-647 Halo^32^ in 100 ul in PBS at RT. Next, the sample was washed three times with PBS, and incubated for 15 minutes on a Poly-L-lysine-coated coverslip. The excess sample was removed after which the slide is mounted on a microscope slide using Mowiol. The sample slides were stored at 4 degrees Celsius in the dark until use.

### Software

The software to control the Abberiort Expert Line STED microscope is Imspector v16.3.11069-w2020 (https://imspector.abberior-instruments.com/). Imspector allows for controlling the microscope through Python code through the package Specpy v1.2.3 (https://pypi.org/project/specpy/). Python 3.7.9 is used to develop the code for the presented automation.

## Data availability

The data that support the findings of this study are available from the corresponding author, R.V., upon reasonable request.

## Code availability

The code that support the findings of this study are available from the corresponding author, R.V., upon reasonable request.

## Author contributions

R. V. designed and supervised the research. R.V. and F.M. developed the code to automate STED measurements. F.M. and R.V. wrote the manuscript.

## Acknowledgement

We thank Thomas C.Q. Burgers and Anna Kenbeek (Zernike Institute for Advanced Materials, Rijksuniversiteit Groningen) for initial work on the Python communication with the microscope. We also thank dr. Fredrik Hurtig and dr. Gabriel Tarrason-Risa for preparation of the Archaea *Sulfolobus acidocaldarius* sample slides, and Prof. dr. Buzz Baum and dr. Fredrik Hurtig for fruitful discussions on the requirements for optimal imaging and analysis of the Archaea *S. acidocaldarius* samples (University College London). Furthermore, we thank Prof. dr. Ida J. van der Klei and Eline M.F. de Lange for the kind gift of the yeast sample, and Alexey N. Butkevich (Max Planck Institute for Medical Research) for the SiR647-Halo dye used in this sample. We thank Prof. dr. Buzz Baum and dr. Jessica Matthias for careful reading of the manuscript.

---

**Supplementary 1.**
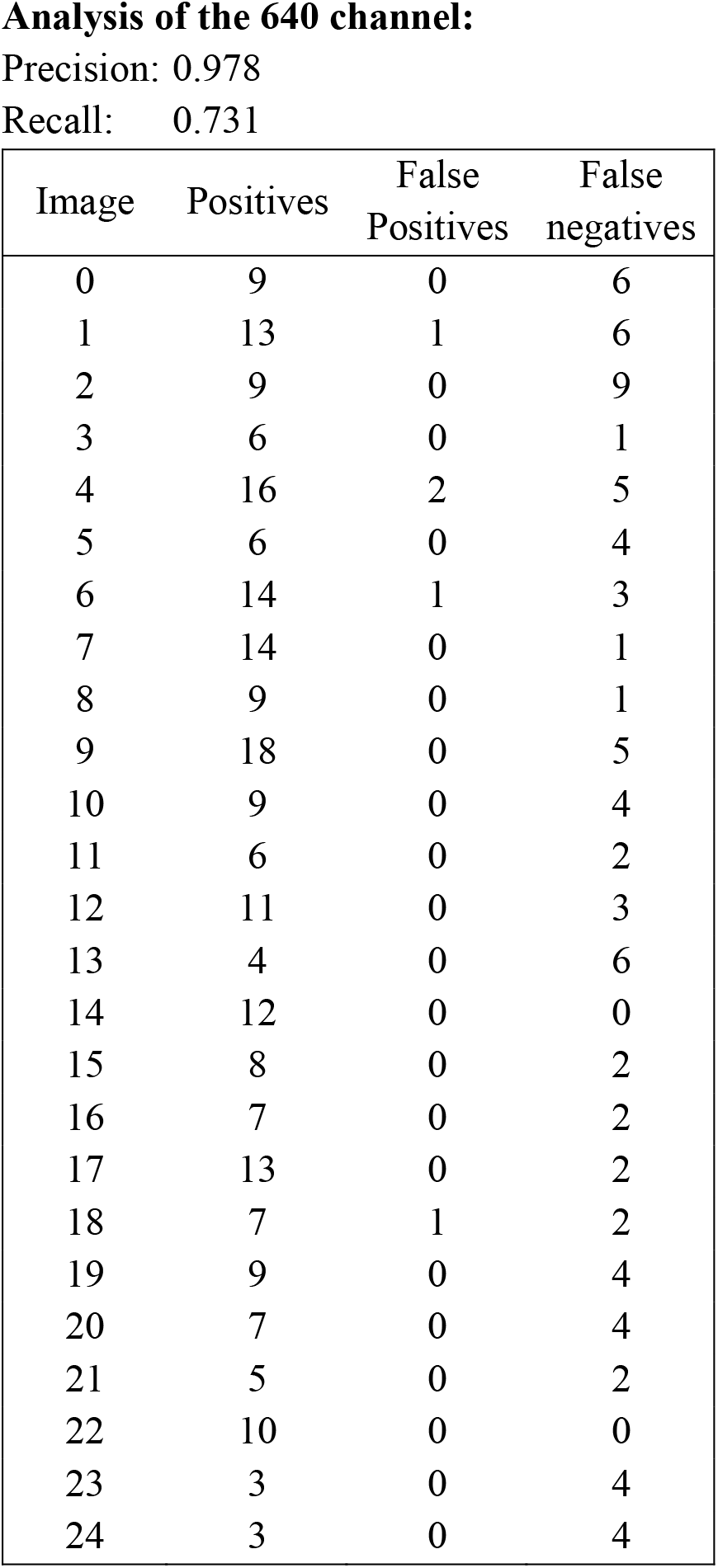
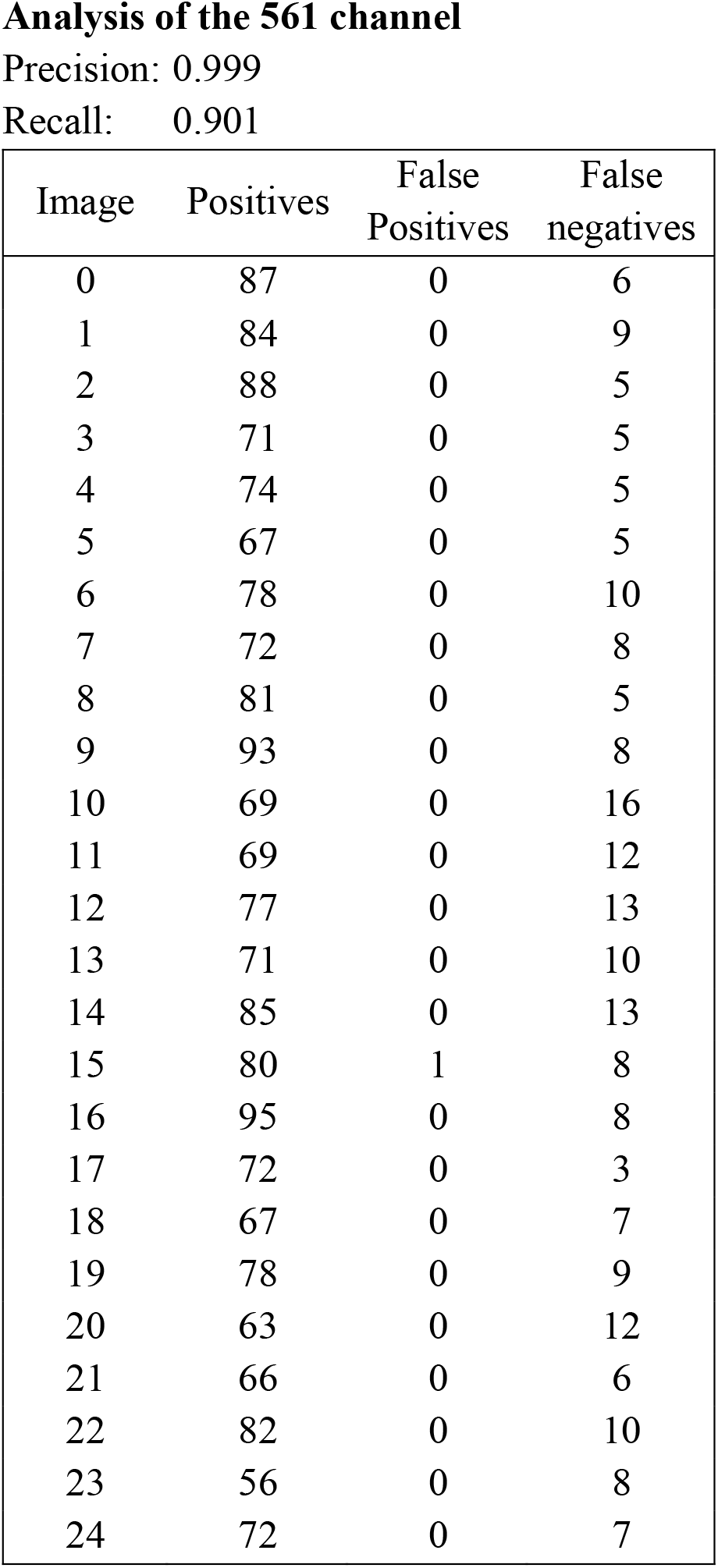
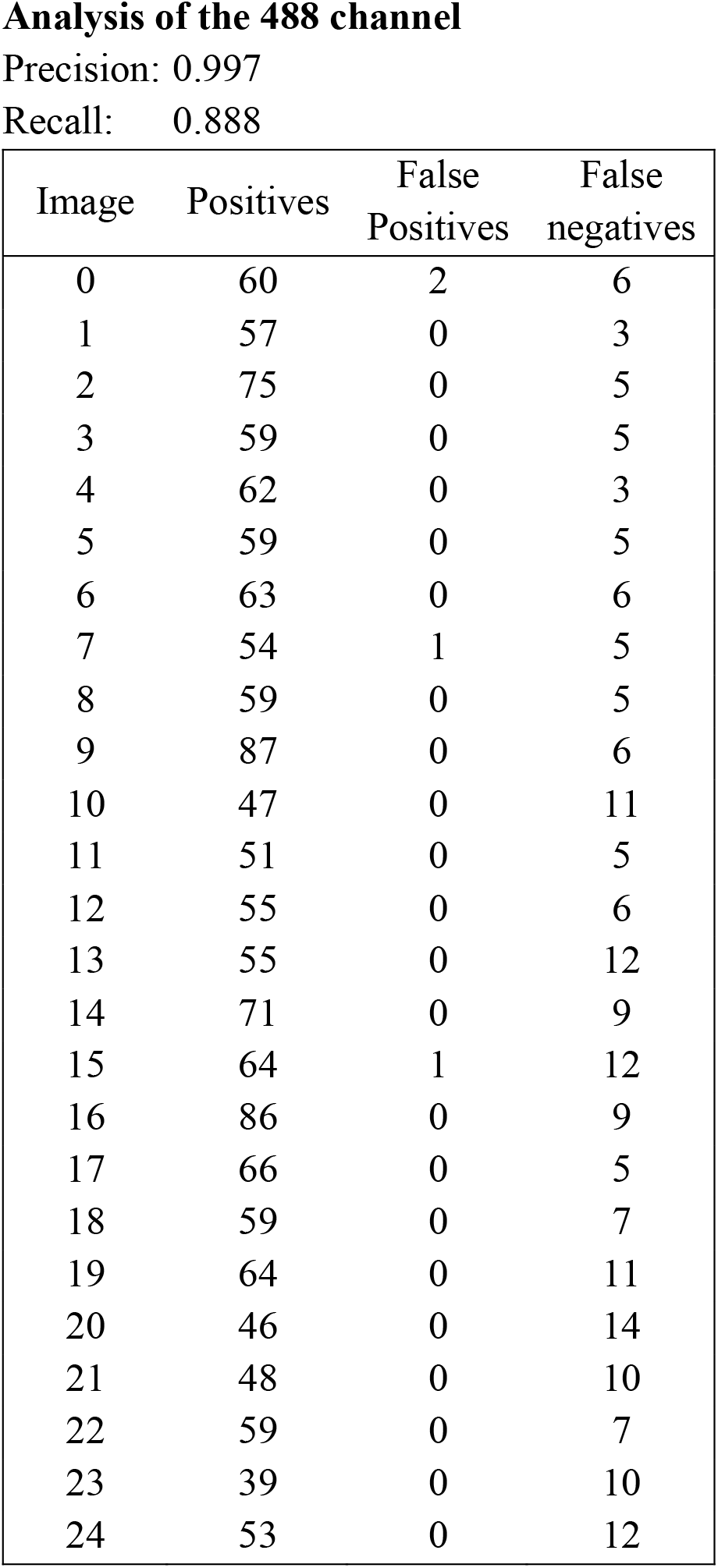
Segmentation results of 25 overview images with DNA (405 channel), and three cell division proteins (640, 561, 488 excitation channels)

| Image | Positives | False Positives | False negatives |
| --- | --- | --- | --- |
| 0 | 9 | 0 | 6 |
| 1 | 13 | 1 | 6 |
| 2 | 9 | 0 | 9 |
| 3 | 6 | 0 | 1 |
| 4 | 16 | 2 | 5 |
| 5 | 6 | 0 | 4 |
| 6 | 14 | 1 | 3 |
| 7 | 14 | 0 | 1 |
| 8 | 9 | 0 | 1 |
| 9 | 18 | 0 | 5 |
| 10 | 9 | 0 | 4 |
| 11 | 6 | 0 | 2 |
| 12 | 11 | 0 | 3 |
| 13 | 4 | 0 | 6 |
| 14 | 12 | 0 | 0 |
| 15 | 8 | 0 | 2 |
| 16 | 7 | 0 | 2 |
| 17 | 13 | 0 | 2 |
| 18 | 7 | 1 | 2 |
| 19 | 9 | 0 | 4 |
| 20 | 7 | 0 | 4 |
| 21 | 5 | 0 | 2 |
| 22 | 10 | 0 | 0 |
| 23 | 3 | 0 | 4 |
| 24 | 3 | 0 | 4 |

| Image | Positives | False<br>Positives | False<br>negatives |
| --- | --- | --- | --- |
| 0 | 87 | 0 | 6 |
| 1 | 84 | 0 | 9 |
| 2 | 88 | 0 | 5 |
| 3 | 71 | 0 | 5 |
| 4 | 74 | 0 | 5 |
| 5 | 67 | 0 | 5 |
| 6 | 78 | 0 | 10 |
| 7 | 72 | 0 | 8 |
| 8 | 81 | 0 | 5 |
| 9 | 93 | 0 | 8 |
| 10 | 69 | 0 | 16 |
| 11 | 69 | 0 | 12 |
| 12 | 77 | 0 | 13 |
| 13 | 71 | 0 | 10 |
| 14 | 85 | 0 | 13 |
| 15 | 80 | 1 | 8 |
| 16 | 95 | 0 | 8 |
| 17 | 72 | 0 | 3 |
| 18 | 67 | 0 | 7 |
| 19 | 78 | 0 | 9 |
| 20 | 63 | 0 | 12 |
| 21 | 66 | 0 | 6 |
| 22 | 82 | 0 | 10 |
| 23 | 56 | 0 | 8 |
| 24 | 72 | 0 | 7 |

| Image | Positives | False<br>Positives | False<br>negatives |
| --- | --- | --- | --- |
| 0 | 60 | 2 | 6 |
| 1 | 57 | 0 | 3 |
| 2 | 75 | 0 | 5 |
| 3 | 59 | 0 | 5 |
| 4 | 62 | 0 | 3 |
| 5 | 59 | 0 | 5 |
| 6 | 63 | 0 | 6 |
| 7 | 54 | 1 | 5 |
| 8 | 59 | 0 | 5 |
| 9 | 87 | 0 | 6 |
| 10 | 47 | 0 | 11 |
| 11 | 51 | 0 | 5 |
| 12 | 55 | 0 | 6 |
| 13 | 55 | 0 | 12 |
| 14 | 71 | 0 | 9 |
| 15 | 64 | 1 | 12 |
| 16 | 86 | 0 | 9 |
| 17 | 66 | 0 | 5 |
| 18 | 59 | 0 | 7 |
| 19 | 64 | 0 | 11 |
| 20 | 46 | 0 | 14 |
| 21 | 48 | 0 | 10 |
| 22 | 59 | 0 | 7 |
| 23 | 39 | 0 | 10 |
| 24 | 53 | 0 | 12 |

## Supplementary figures

**Supplementary Figure 1.**
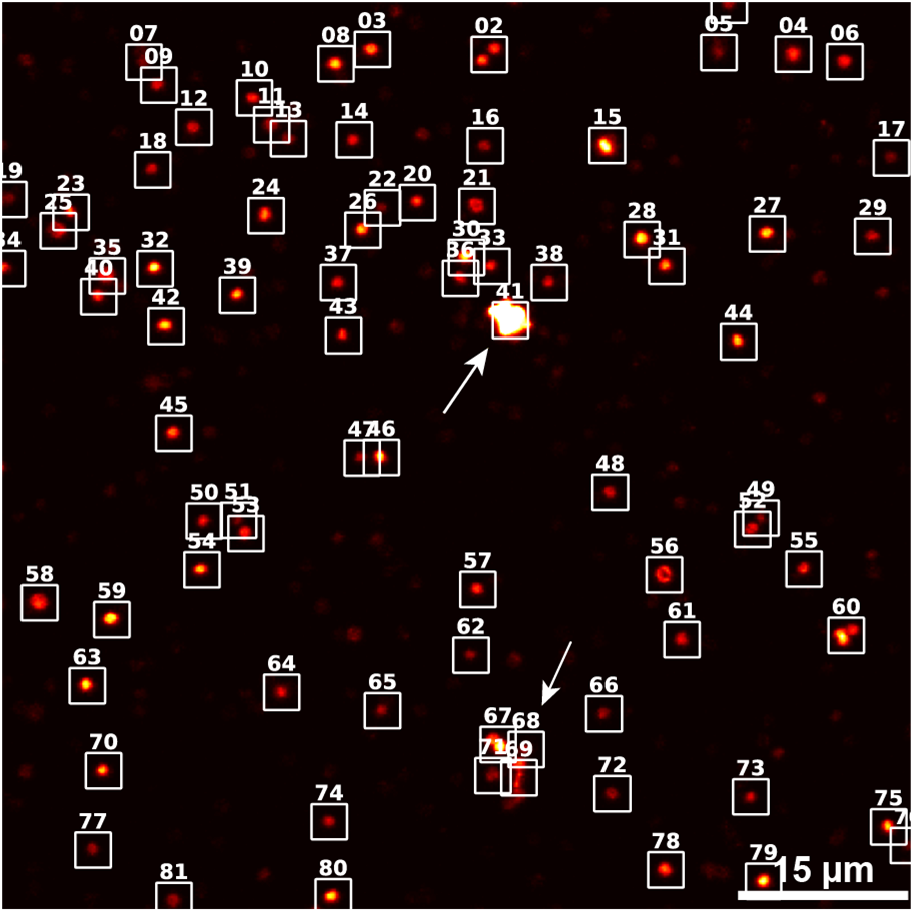
*Segmentation map* of a CdvB2 labelled with Alexa Fluor 488 and imaged in the 488nm excitation channel. The segmentation shows two falsely positive segmented objects (arrows). Both objects show noise. FOV is 80×80µm^2^.

**Supplementary Figure 2.**
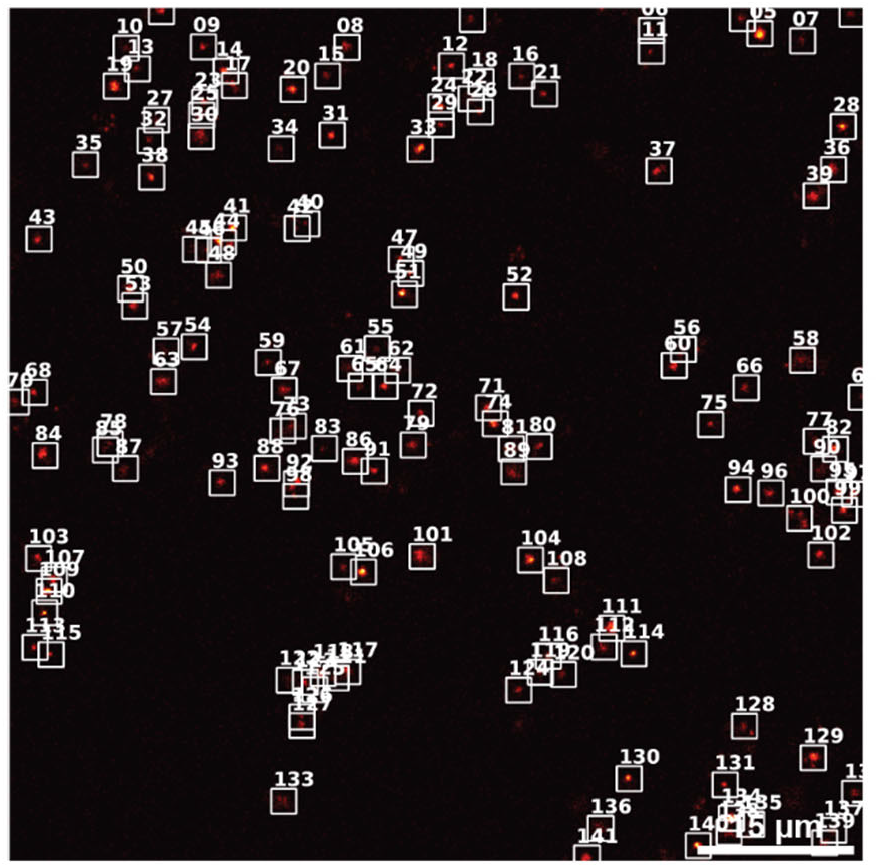
*Segmentation map* of a single FOV (80×80µm^2^) of a peroxisomal protein labelled in *Hansenula polymorpha* (yeast). Using our developed automation method, a dataset of 8,063 STED images of individually measured cells had been acquired.

**Supplementary Figure 3.**
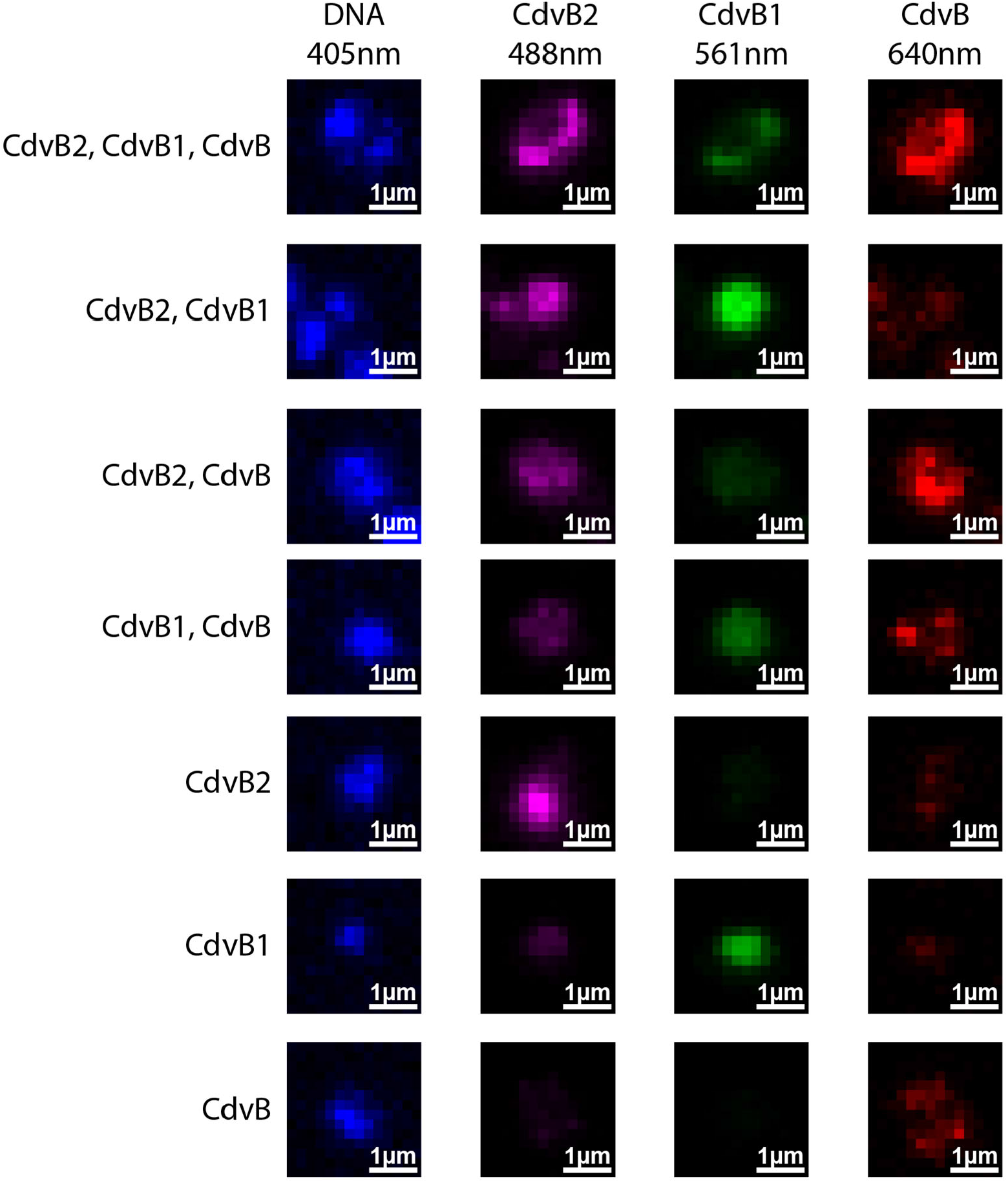
Examples of the identified combinations of protein presence of ESCRT-III proteins involved in cell division in archaea. The combination of present proteins in close proximity is determined in the selection step, using the segmented coordinates from the overview images. The above shown examples show all the different combinations of present ESCRT-III proteins found in the sample. Here, CdvB2 was labelled with Alexa Fluor 488 (magenta, 488nm channel), CdvB1 was labelled with Abberior STAR 580 (green, 561nm channel) and CdvB was labelled with Abberior STAR 635 (red, 640nm channel). Additionally, DNA (blue, 405nm channel) was imaged. Each row is a representation of one specific subregion from the sample. The pixel size in each image is 200×200nm^2^, and the regions displayed are 2.8×2.8µm^2^.

